# ZFP423 regulates early patterning and multiciliogenesis in the hindbrain choroid plexus

**DOI:** 10.1101/2020.03.04.975573

**Authors:** Filippo Casoni, Laura Croci, Francesca Vincenti, Paola Podini, Luca Massimino, Ottavio Cremona, G. Giacomo Consalez

## Abstract

The choroid plexus (ChP) is a secretory tissue that produces cerebrospinal fluid (CSF) and secretes it into the ventricular system. CSF flows from the lateral to the third ventricle, and then to the fourth ventricle through the cerebral aqueduct. Recent studies have uncovered new, active roles for this structure in the regulation of neural stem cell maintenance and differentiation into neurons. *Zfp423,* encoding a Kruppel-type zinc finger transcription factor essential for cerebellar development and mutated in rare cases of cerebellar vermis hypoplasia / Joubert syndrome and other ciliopathies, is expressed in the hindbrain roof plate (RP), from which the IV ventricle ChP arises, and in mesenchymal cells giving rise to the stroma and leptomeninges. *Zfp423* mutants display a marked reduction of the hindbrain ChP (hChP), which fails to express key markers of its secretory function and genes implicated in its development and maintenance (*Lmx1a, Otx2*). The mutant hChP displays a complete lack of multiciliated ependymal cells. A transcriptome analysis conducted at the earliest stages of hChP development and subsequent validations demonstrate that the mutant hChp displays a strong deregulation of pathways involved in early hindbrain patterning and multiciliated cell fate specification. Our results propose *Zfp423* as a master gene and one of the earliest known determinants of hChP development.

## INTRODUCTION

The choroid plexus (ChP) is a secretory structure consisting of an epithelial cell layer vascularized by fenestrated blood vessels. The ChP produces cerebrospinal fluid (CSF) and releases it into all ventricles of the vertebrate brain, from which the CSF flows into the central canal of the spinal cord and into the subarachnoid space (Damkier et al.,2013; Lehtinen and Walsh, 2011). The CSF plays an essential role in the CNS (inter alia Lehtinen et al., 2013) and contains multiple signals, including growth factors (Parada et al., 2006; Zappaterra et al., 2007), necessary for neural development. Conversely, CSF overproduction, obstructed flow or inadequate resorption lead to hydrocephalus (Damkier et al., 2013). Philogenetically, the ChP and its function are conserved from lower vertebrates to humans (Brocklehurst, 1979).

The ChP develops on the dorsal aspect of the neural tube at three locations: fourth, third and lateral ventricles, in the hindbrain, diencephalon and telencephalon, respectively. The hindbrain ChP (hChP) of the fourth ventricle appears first, followed by the appearance of the telencephalic ChP (tChP) in the lateral ventricles, and finally by the diencephalic ChP in the third ventricle (reviewed in Lehtinen et al., 2013). The secretory role of the ChP reflects its structure that consists of a monolayer of cuboidal epithelial cells, which undergo a series of morphological transformations, molecularly and temporally regulated during development (inter alia Awatramani et al., 2003). ChP epithelial cells, a specialized form of ependyma, derive from neuroepithelial progenitors. At first, they are pseudostratified, then become columnar, and eventually monostratified and cuboidal (reviewed in Lehtinen et al., 2013), and are characterized by the presence of tufts of cilia and microvilli. Transition through these distinct stages is accompanied by structural changes such as epithelial polarization and formation of numerous microvilli and several cilia along the apical domain of the cell membrane (Sturrock, 1979). While mature ChP epithelia are histologically and ultrastructurally indistinguishable, irrespective of location, tChP and hChP are heterogeneous in regard to their transcriptional profiles and positional identities, and they are also functionally distinct, secreting two region-specific varieties of CSF (reviewed in Lehtinen et al., 2013).

Hindbrain ChP specification occurs between embryonic day 8.5 (E8.5) and E9.5 (Thomas and Dziadek, 1993), 2–3 days before hChP morphology unveils itself. The hChP epithelium derives from neuroectodermal roof plate progenitors positive for the bone morphogenetic protein (BMP)-family ligand GDF7 (Currle et al., 2005), while the stromal component is comprised of mesenchymal cells of mesodermal origin (Wilting and Christ, 1989). The adoption of ChP cell fate is established early in development and requires the expression of Notch signaling targets *Hes1, Hes3* and *Hes5* (Imayoshi et al., 2008). LIM-homeobox protein Lmx1a is expressed in the hindbrain rhombic lip (Chizhikov et al., 2010). In *Lmx1a*-/- (*Dreher*) mice, the hindbrain roof plate does not form, leading to a failure of hChP development (Millonig et al., 2000). Likewise, conditional *Otx2* deletion in E9 embryos impairs the development of all ChPs, while its abrogation at a later stage (E15) affects only the hChP (Johansson et al., 2013). The role played at later stages (E12.5-E14.0) by the lower rhombic lip (LRL) in hChP progenitor proliferation has been elegantly described: early-born, RP-derived hChP ependymal cells differentiate, exit the cell cycle, and start releasing the mitogen sonic hedgehog (SHH), which promotes the expansion of a mitotic progenitor pool located in the transition zone, adjacent to the LRL (Huang et al., 2009b). Conversely, the role played in hChP development by the upper rhombic lip (URL), if any, has not been clarified. More broadly, the earliest molecular mechanisms leading to hChP formation are incompletely characterized.

The *Zfp423* gene and its human homolog *ZNF423* encode a 30-zinc-finger transcription factor involved in key developmental pathways. ZFP423 acts both as a transcription factor and as a scaffold for multiple protein-protein interactions with signaling molecules involved in early patterning, cell fate specification, neurogenesis and various aspects of differentiation (Casoni et al., 2017; Hata et al., 2000; Hong and Hamilton, 2016; Huang et al., 2009a; Masserdotti et al., 2010; Massimino et al., 2018; Tsai and Reed, 1998). Moreover, ZFP423 has been implicated in the DNA damage response in vitro (Chaki et al., 2012; Ku et al., 2003) and in vivo (Casoni et al., 2017). While all homozygous mutants described to date exhibit severe cerebellar malformations (Alcaraz et al., 2006; Casoni et al., 2017; Cheng et al., 2007; Warming et al., 2006), two allelic mutant lines described by our group, in each of which a distinct functional domain is deleted, have revealed domain-specific functions in cerebellar ventricular zone neurogenesis (Casoni et al., 2017). ZFP423, which is required for proper cerebellar granule cell development, also plays an important role in ciliary development/function in mitotic monociliated neurogenic progenitors of the external granular layer (Hong and Hamilton, 2016). In humans, *ZNF423* mutations have been linked to cerebellar vermis hypoplasia, Joubert syndrome 19 and nephronophthisis 14 (Chaki et al., 2012).

Previous studies have reported a malformation of the choroid plexus in spontaneous *Zfp423* mutants (Alcaraz et al., 2006). In the present paper we report the expression of the ZFP423 protein in the roof plate and developing hChP. Moreover, by analyzing a C-terminal *Zfp423* mutant that displays a near-complete deletion of this structure, we examine the roles played by the homonymous protein at the earliest stages of hChP morphogenesis, and its contribution to the development of mesenchymal and epithelial components of this tissue. Although this phenotype is present in both allelic mutants (Casoni et al., 2017), we focused our analysis on the Δ28-30 line.

## MATERIALS AND METHODS

### Animal Care

All experiments described in this paper were performed in agreement with the stipulations of the San Raffaele Scientific Institute Animal Care and Use Committee (I.A.C.U.C.).

### Mouse genetics

The *Zfp423* Δ28-30 mouse lines were generated by gene targeting as previously described (Casoni et al., 2017). All experiments were performed using homozygous mutant animals starting from backcross generation N10, which carry 99,8% C57BL/6N genetic background. All studies were conducted in homozygous mutant embryos, using co-isogenic wild type control littermates. Genotyping was done by PCR as described (Corradi et al., 2003) using allele-specific primers. Primer sequences are listed under Supplemental Materials and Methods.

### Tissue Preparation

Embryos were collected from time-mated dams and the emergence of the copulation plug was taken as E0.5. For the preparation of embryonic samples, pregnant dams were anesthetized with Avertin (Sigma). Embryos were fixed 6-8 hours according to their developmental stage by immersion with 4% PFA in 1X PBS, cryoprotected overnight in 30% sucrose in 1X PBS, embedded in OCT (Bioptica), and stored at −80°C. For the preparation of postnatal brains, mice were anesthetized with Avertin (Sigma), transcardially perfused with 0.9% NaCl, followed by 4% PFA in 1X PBS, and prepared for freezing as above. The brains and embryos were sectioned sagittally or frontally on a cryotome.

Samples were cryosectioned using CM3050S Leica cryostat and collected on Superfrost plus slides (VWR). Embryonic sections thickness was 14 μm for immunostainings and in situ hybridizations (ISH); Embryonic sections thickness was 30 μm.

### Immunofluorescence

Sections were washed in 1X PBS, blocked and permeabilized in 10% serum, 0.3% Triton X-100 in 1X PBS, and incubated with primary antibodies overnight at 4°C, rinsed, then incubated with species-appropriate secondary antibodies at room temperature for 2 hrs (1:1000: Alexa Fluor-488, Alexa Fluor-568, or Alexa Fluor-647; Molecular Probes^®^). Sections were counterstained with DAPI (1:5000, Sigma) and mounted with fluorescent mounting medium (Dako).

### Antibodies

Mouse monoclonal antibodies included: ZO-1 (1:100, BD Biosciences); Arl13b H3 (1:5, NeuroMab). Rabbit antibodies included: ZFP423/OAZ XL (1:2000, Santa Cruz Biotechnology); Lmx1a (1:3000, Millipore); Collagen type IV (1:500, NovusBio); Desmin (1:200, Cell Sciences); γTubulin (1:800, Sigma); Ezrin (Upstate, 1:50). Rat antibody included: Pecam 1 (1:200, BD Biosciences). Signals from anti-OAZ and Lmx1a antibodies were amplified using the Tyramide Signal Amplification Kit (Perkin Elmer), according to manufacturer’s instructions. Antigen retrieval was performed in a citrate buffer before incubation with anti Lmx1a antibody.

### In situ hybridization

Digoxigenin-labeled riboprobes were transcribed from plasmids containing *Ttr, Lmx1a, Otx2, Gmnc, GDF7* cDNAs. In situ hybridization was performed as described (Croci et al., 2011).

### RT-qPCR

Total RNA was extracted with RNeasy MiniKit (Qiagen), according to manufacturer’s instructions. 1–1.5 μg of total RNA was retrotranscribed using first strand cDNA MMLV-Retrotranscriptase (Invitrogen) and random primers. Each cDNA was diluted 1:10, and 3 μl were used for each RT-qPCR reaction. mRNA quantitation was performed with SYBR Green I Master Mix (Roche) on a LightCycler480 instrument (Roche). Each gene was analyzed in triplicate, and each experiment was repeated at least three times. Data analysis was performed with the 2-ΔΔC(T) method. All RNA levels were normalized based upon *β-actin, H3f3a and Gapdh* transcript levels. For details see Supplementary Materials and Methods.

### Microscopy and image processing

Confocal microscopy pictures were taken on a Leica SP8 microscope. The objective used to obtaiend the images were the following: 1) 20X: HC PL APO CS2 20X (NA 0.75) Dry; 2) 40x: HC PL APO CS 2 40X (NA 1.3) Oil; 3) 63x: HC PL APO CS 2 63X (NA 1.4) Oil. Epifluorescence pictures were taken on Zeiss Axioplan2 microscope. Images were further analyzed using FIJ/ImageJ software. Each staining was replicated on at least three different animals for each genotype. Figures of the paper were prepared using Adobe Photoshop and only contrast and brightness of the pictures were adjusted.

### Statistical analysis

Statistical analysis was conducted using two-tailed unequal variance t test (Welch t-test) for comparison of sets of quantitative values (wt vs. each mutant), or 2way ANOVA for multiple comparisons. Data were reported as the mean ± SD or s.e.m, as indicated.

### RNA-sequencing

E9.5 hindbrain spanning rhombomeres 1-6 were dissected from wild type and mutant embryos (n=3) to obtain total mRNA preparations (miRNeasy Micro Kit, Qiagen). Sequencing libraries were prepared following the SMART-Seq v4 Ultra Low Input RNA (TaKaRa) protocol. Briefly, 1ng of total RNA was used for cDNA synthesis, followed by PCR amplification. FastQC software was used to assess the quality of the FASTQ files. Reads were aligned to the mouse genome, version mm10, using STAR_2.5.3 (Dobin et al., 2013) and the quality of the alignment assessed with Bamtools (Barnett et al., 2011). The Rsubread package (Liao et al., 2019) was used to assign read counts to the genes of the GENCODE model, version M13 [https://www.gencodegenes.org], with all isoforms of each gene considered together. Genes with at least 1 cpm (count per million) in at least 3 samples were considered expressed and assessed for differential expression. Differentially expressed genes were identified using limma (Ritchie et al., 2015). Genes with nominal pvalue < 0.01 and absolute value of log2 fold change > 1 were considered differentially expressed (Consortium, 2014). Geneset functional enrichment was performed with DAVID (PMID 22543366). Downstream statistics and Plot drawing were performed within the R environment. Heatmaps were generated with GENE-E (GENE-E. Cambridge (MA): The Broad Institute of MIT and Harvard).

## RESULTS

In order to study the role of *Zfp423* in ChP development, we analyzed a previously described mouse line (Casoni et al., 2017) that produces a C-terminally deleted protein. In this line, called *Zfp423^Δ28-30^,* nucleotides 3861-4108 of the ORF have been removed, causing a frameshift mutation that produces a stop codon 158 bp into exon 8. The truncated protein misses 91 C-terminal amino acids, including zinc finger 28-30.

The analysis of Nissl-stained P18 cerebellar frontal sections revealed the presence in wt embryos of an elaborate hChP in the 4th ventricle, which was completely absent in the mutant (Fig. 1A), except for a very small segment of ChP tissue at the very lateral edge of the ventricle (Fig. 1A’, inset). This observation prompted further studies of hChP development. Parasagittal sections were analyzed at earlier ontogenetic stages. At E14.5, mutants displayed an enlarged 4^th^ ventricle and a rudimentary hChP primordium (Fig. 1B’, red arrowhead) when compared to wt littermates (Fig. 1B, white arrowhead). However, in parasagittal sections of the telencephalon, the primordium of the telencephalic ChP (tChP) shows no significant alterations (Fig. 1C,C’, white arrowheads), suggesting that *Zfp423* specifically affects hChP development in the 4^th^ ventricle.

**Figure 1.**
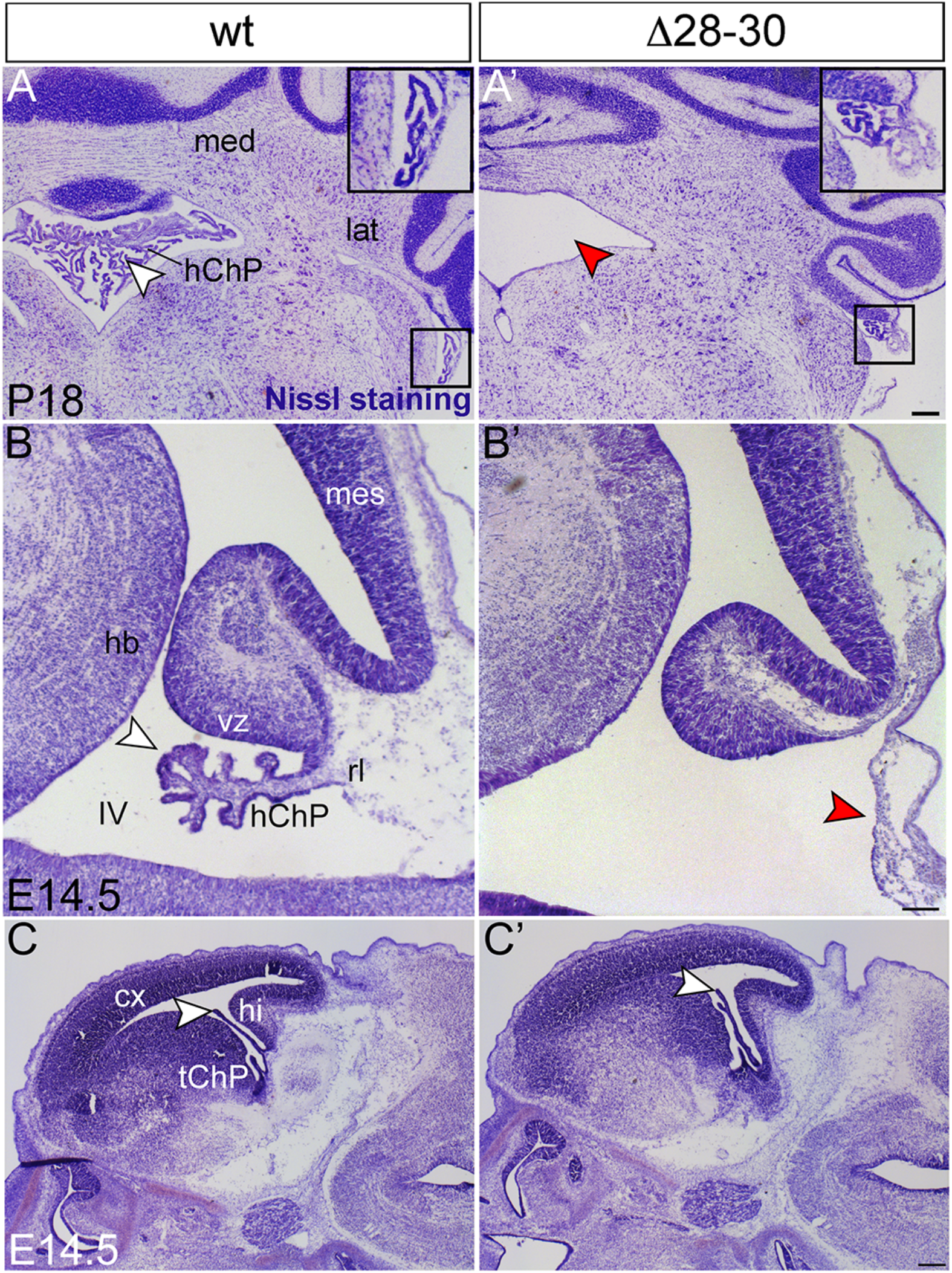
Profound malformation of the mutant hChP. A,A’: Nissl staining of P18 mice frontal sections reveals the near-complete absence of the hindbrain ChP (hChP) in the 4th ventricle of the *Zfp423* mutant (red arrowhead in A’) compared to the wt (A). A small lateral segment of the hChP is visible in wt and mutants (insets). B,B’: Nissl staining of E14.5 parasagittal sections shows a severe hChP hypoplasia in the mutant compared to wt, with a flattened surface (arrowhead in B’). C,C’: Conversely, the mutant’s tChP shows a rather normal morphology compared to the wt. med: medial cerebellar nucleus; lat: lateral nucleus. cx: cortex; hChP: hindbrain choroid plexus; tChP: telencephalic choroid plexus; hb: ventral hindbrain; hi: hippocampus; IV: fourth ventricle; mes: mesencephalon; rl: rhombic lip; vz: ventricular zone. Size bar: 200μm in A, 100μm in B, 400μm in C.

Transthyretin (Ttr) is one of the earliest functional markers of the ChP: it is a protein that transports the thyroid hormone thyroxine (T4) and retinol-binding protein bound to retinol (Power et al., 2000; Raz and Goodman, 1969) from ChP epithelial cells into the CSF. *In situ* hybridization was used to analyze *Ttr* expression in the E11.5 hChP. Our results indicate that the expression of the *Ttr* gene is dramatically reduced in the mutant (Fig. 2A’,B’, red arrowheads) compared to controls (Fig. 2A,B, black arrowheads). Moreover, by quantitative RT-PCR (RT-qPCR), we analyzed mRNA levels of *Ttr* and *aquaporin 1 (Aqp1),* which encodes a water transporter highly expressed in the ChP. Our results confirmed a sharp downregulation of *Ttr* in mutants at E11.5 (*** p< 0.0001, Fig. 2C) and E13.5 (*** p< 0.0001, Fig. 2D). Similarly, *Aqp1* mRNA was downregulated in mutants at E13.5 (**** p < 0.00001, Fig. 2E). Thus, *Zfp423* mutation affects both hChP morphology and function during embryonic development.

**Figure 2.**
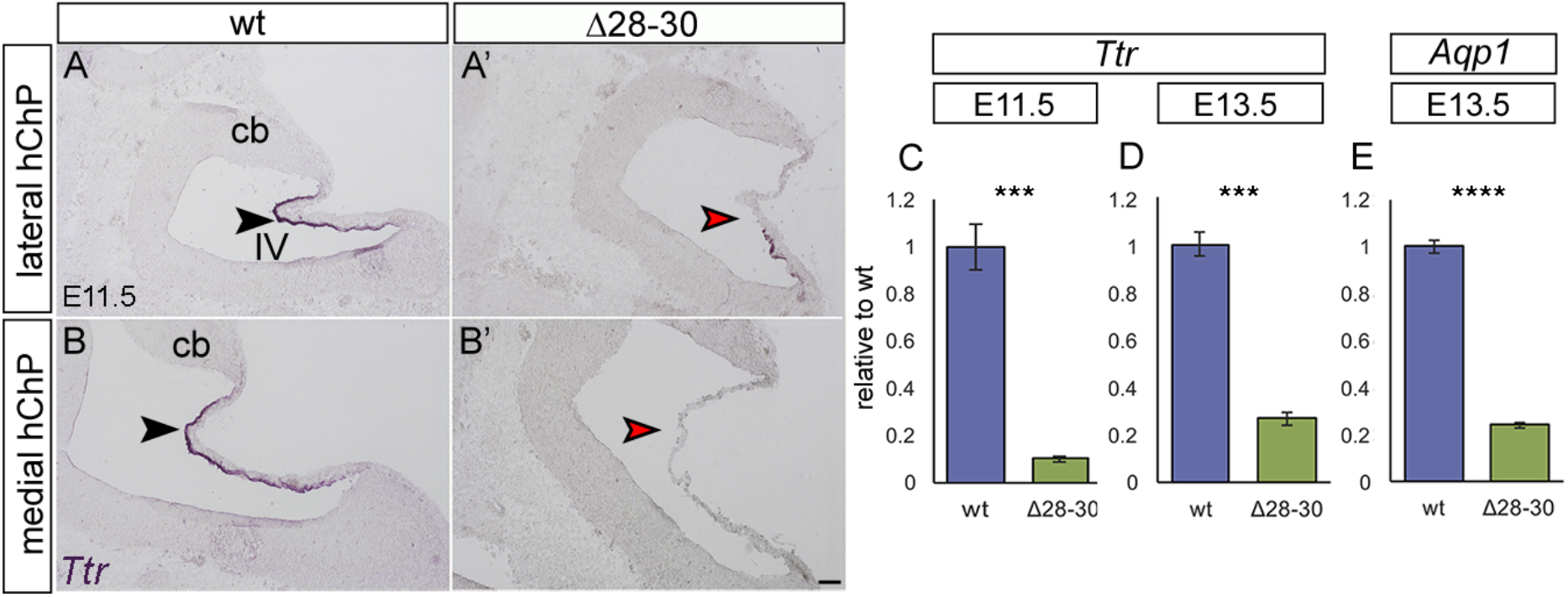
Functional markers of the hChP are downregulated in the *Zfp423-mutant* hChP during development. A-B’: *in situ* hybridization of parasagittal E11.5 sections, lateral and medial, done with a *Ttr* riboprobe. *Ttr* is downregulated and the expression is patchy in lateral sections of the mutant hChP epithelium (A’), while it is completely absent in medial parasagittal sections (B’). RT-qPCR reveals a sharp downregulation of *Ttr* in mutants at E11.5 (C, N=3/genotype; and E13.5 (D, N=5/genotype). The *Aqp1* gene is also downregulated in mutants at E13.5 (E, N=3/genotype). Data are plotted as mean ± s.e.m; Welch’s unequal variances t-test; **** p< 0.00001, *** p< 0.0001. cb: cerebellum; IV: fourth ventricle. Size bar: 100μm in A-B’.

### ZFP423 protein expression

To define the possible role of *Zfp423* in hChP development, we first investigated the precise distribution of the corresponding protein in the hChP. We immunostained sagittal rhombencephalic sections with a previously characterized anti ZFP423 polyclonal Ab (Casoni et al., 2017; Hong and Hamilton, 2016; Shao et al., 2016). At E11.5, ZFP423 was highly expressed in the developing hChP epithelium (Fig. 3A, white arrowheads), in the upper and lower rhombic lip (URL and LRL, respectively) (Fig. 3A, white arrows). At E12.5, ZFP423 remains strongly expressed in different areas, namely URL and LRL (Fig. 3B, white arrows), hChP mesenchyme and prospective meninges (Fig. 3B, red arrows), and in the caudal third of the hChP epithelium (transition zone, Fig. 3B,B’, arrowheads), while in the rostral two thirds of the hChP epithelium ZFP423 protein expression is no longer detectable.

**Figure 3.**
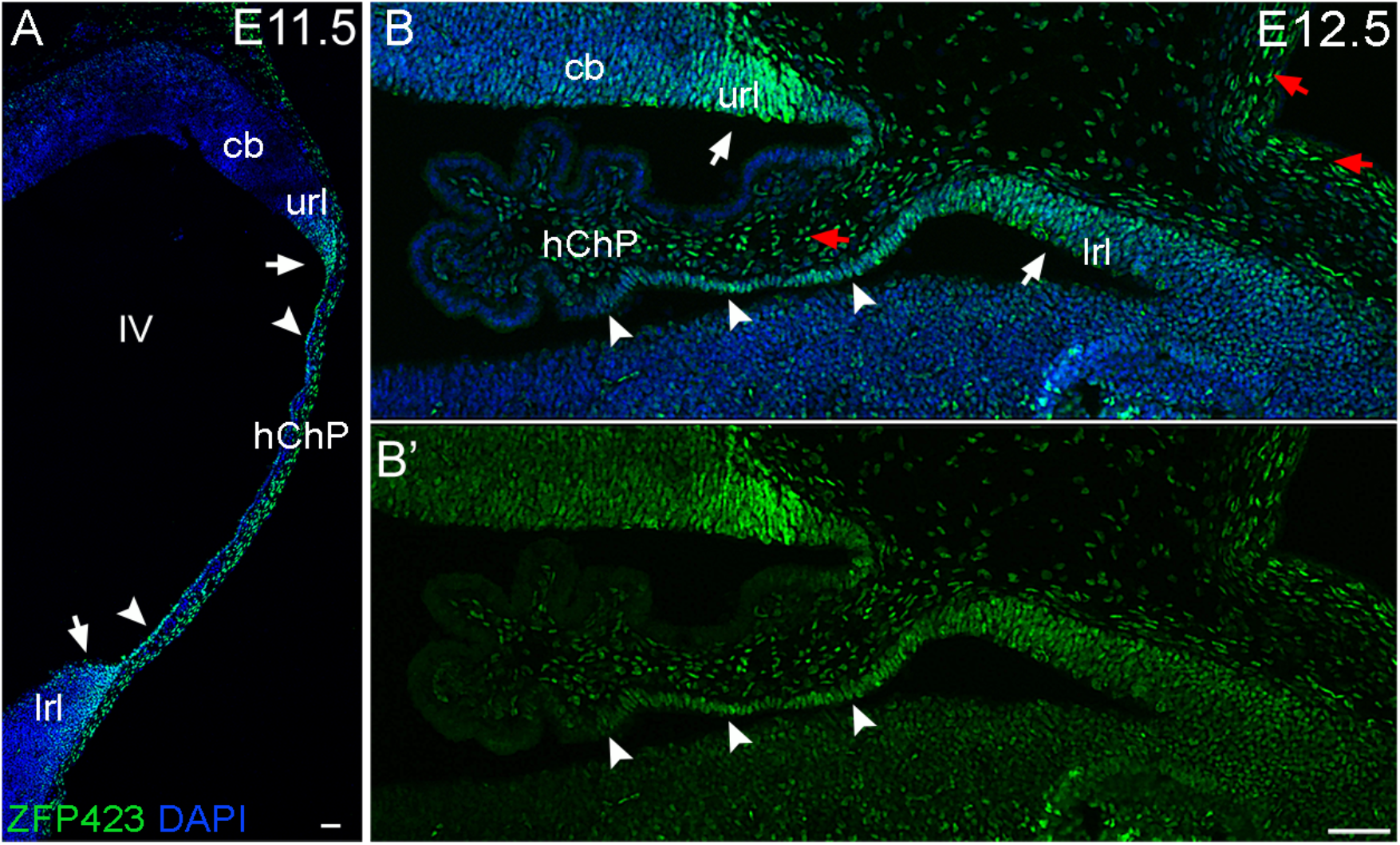
ZFP423 is expressed in ectodermal and mesodermal derivatives of the developing hChP. Immunostaining of parasagittal cerebellar sections at E11.5 (A) and E12.5 (B,B’) reveals ZFP423 protein distribution in the upper and lower rhombic (white arrows), in the developing roof plate epithelium (prospective hChP, white arrowheads), and in the prospective leptomeninges (red arrows). hChP: hindbrain choroid plexus; cb: cerebellum; url: upper rhombic lip; lrl: lower rhombic lip; IV: fourth ventricle. Size bar: 50μm in A, 50μm in B,B’.

### Expression of early ChP markers

In an attempt to understand the molecular mechanisms leading to hChP malformation in *Zfp423* mutants, we analyzed the expression of established hChP markers, namely Lmx1a and Otx2 by *in situ* hybridization and RT-qPCR. Lmx1a is a transcription factor that plays an important role in hChP development (Chizhikov et al., 2010). *Otx2* is another key player in the development and maintenance of this territory (Johansson et al., 2013).

By *in situ* hybridization at E13.5, *Lmx1a* and *Otx2* expression (Johansson et al., 2013) was sharply downregulated in the epithelium of the mutant hChP (Fig. 4A’,B’ red arrowheads), as compared to wt littermates (Fig. 4A,B black arrowheads). Accordingly, RT-qPCR conducted on embryonic hindbrain mRNA confirmed downregulation of both markers at E11.5 and E13.5 in the mutant (Fig. 4C,D). In particular, *Lmx1a* transcript was 30% reduced in the homozygous *Zfp423* mutant hindbrain compared to controls at E11.5, and 50% reduced at E13.5 (Fig. 4C, *** p < 0.0001 and ** p <0,005, respectively). *Otx2* transcript was 50% reduced in the mutant hindbrain at E11.5 and E13.5 (Fig. 4D, ** p < 0,005 and * p < 0,05, respectively). This evidence was confirmed by Lmx1a immunostaining of E11.5, E12.5 and E13.5 sagittal sections: at all stages, Lmx1a decorates the epithelium of the wt hChP (Suppl. Fig 1A, white arrowheads) and its levels increase during development (Suppl. Fig 1B-E, white arrowheads). At the same stages, Lmx1a is discontinuously expressed in the mutant hChP epithelium (Suppl. Fig 1A’, red arrowheads mark Lmx1a–negative epithelium). In the mutant, Lmx1a expression is reduced at E12.5 and almost abolished at E13.5 in medial sections (Suppl. Fig 1B’-E’ red arrowheads). Taken together, our findings suggest that *Zfp423* mutants display a severe loss of two genes known to play a critical role in hChP development and maintenance.

**Figure 4.**
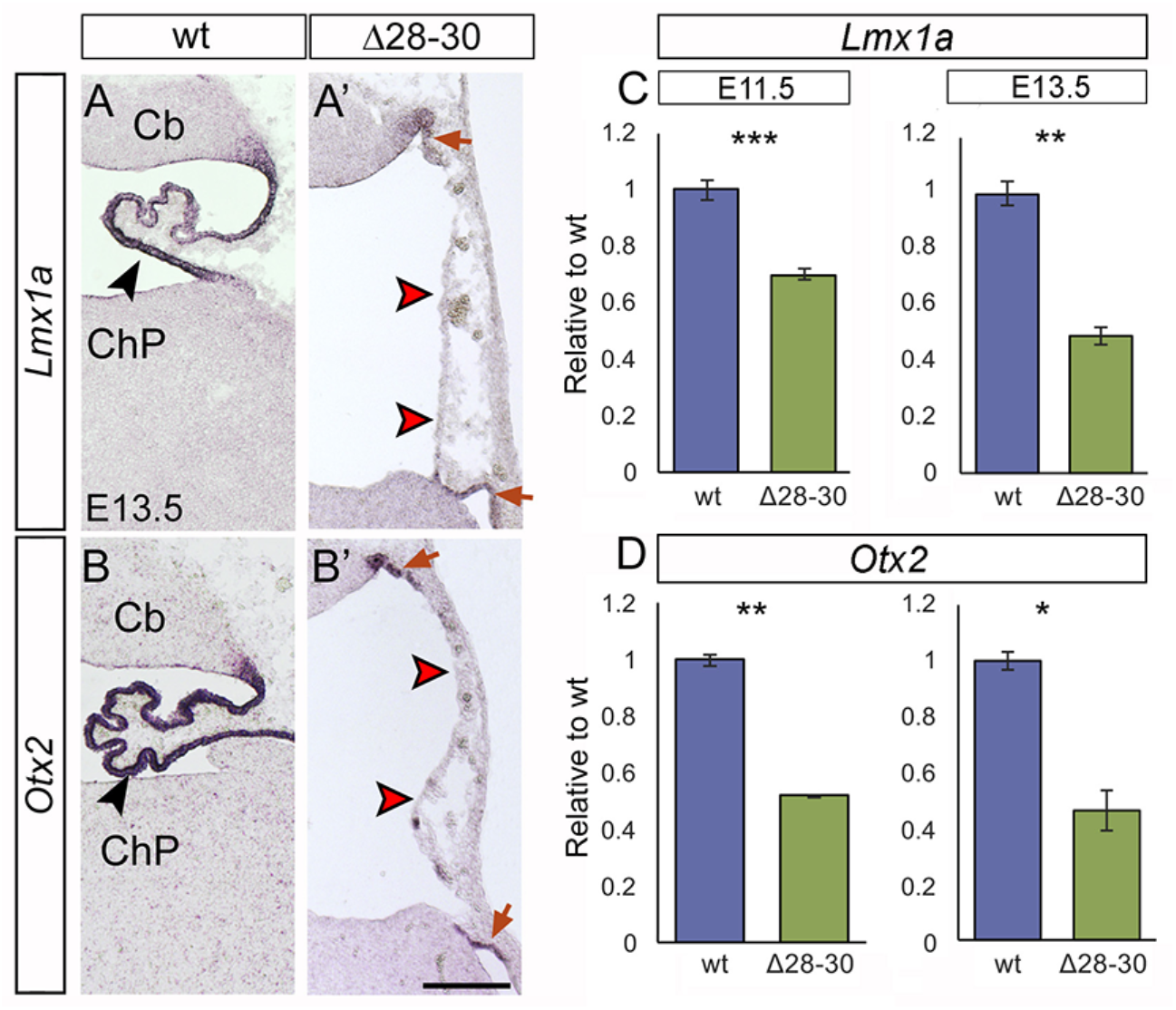
Established markers of the developing hChP are sharply downregulated in the mutant. *In situ* hybridization of cerebellar parasagittal sections at E13.5 with *Lmx1a* and *Otx2* riboprobes. In the mutant hChP, *Lmx1a* (A’) and *Otx2* (B’) are faintly expressed in the upper and lower rhombic lip, which are malformed (red arrows), and absent in the hChP, which is misfolded (red arrowheads). RT-qPCR reveals a sharp downregulation of *Lmx1a* in the mutant hindbrain at E11.5 and E13.5 (C, N=3/genotype; N=5/genotype, respectively) and of *Otx2* in the mutant hindbrain at E11.5 and at E13.5 (D, N= 3/genotype, N= 5/genotype, respectively). Mean ± s.e.m is plotted; Welch’s unequal variances t-test; *** p< 0.0001 and ** p< 0.005, * p< 0.05. cb: cerebellum; hChP: hindbrain choroid plexus. Size bar: 200μm in A-B’.

### Development of the hChP stroma

To begin dissecting the cascade of cell-cell interactions leading to hChP malformation in *Zfp423* mutants, we used immunofluorescence to analyze markers of the hChP stroma. To this end, we immunostained the hChP for (*i*) Platelet Endothelial Cell Adhesion Molecule 1 (PECAM-1), to visualize endothelial cells; (*ii*) Desmin for pericytes, which regulate blood-CSF barrier permeability (Armulik et al., 2010; Daneman et al., 2010); (*iii*) collagen type IV for the lamina densa of the basal membrane secreted by epithelial and endothelial cells; and (*iv*) *Akap12* and *Igf2* for prospective leptomeninges; Our results revealed that (*i*) capillaries are present at E11.5 in the hChP stroma in wt and mutant embryos alike, albeit discontinuous in the latter (Fig. 5A,A’, white arrowheads); (*ii*) Desmin+ pericytes are maintained in mutant embryos at E12.5 (Fig. 5B,B’); (*iii*) collagen type IV is present in the basal membrane of the mutant hChP at E14.5 (Fig. 5C,C’), and (iv) meningeal markers are either mildly downregulated or unchanged in the mutant, as shown by RT-qPCR (Fig. 5D). These findings suggest that the blood supply to the hChP is relatively preserved in *Zfp423* mutants at early stages development, and that the basement membrane is conserved, albeit reduced in its extension.

**Figure 5.**
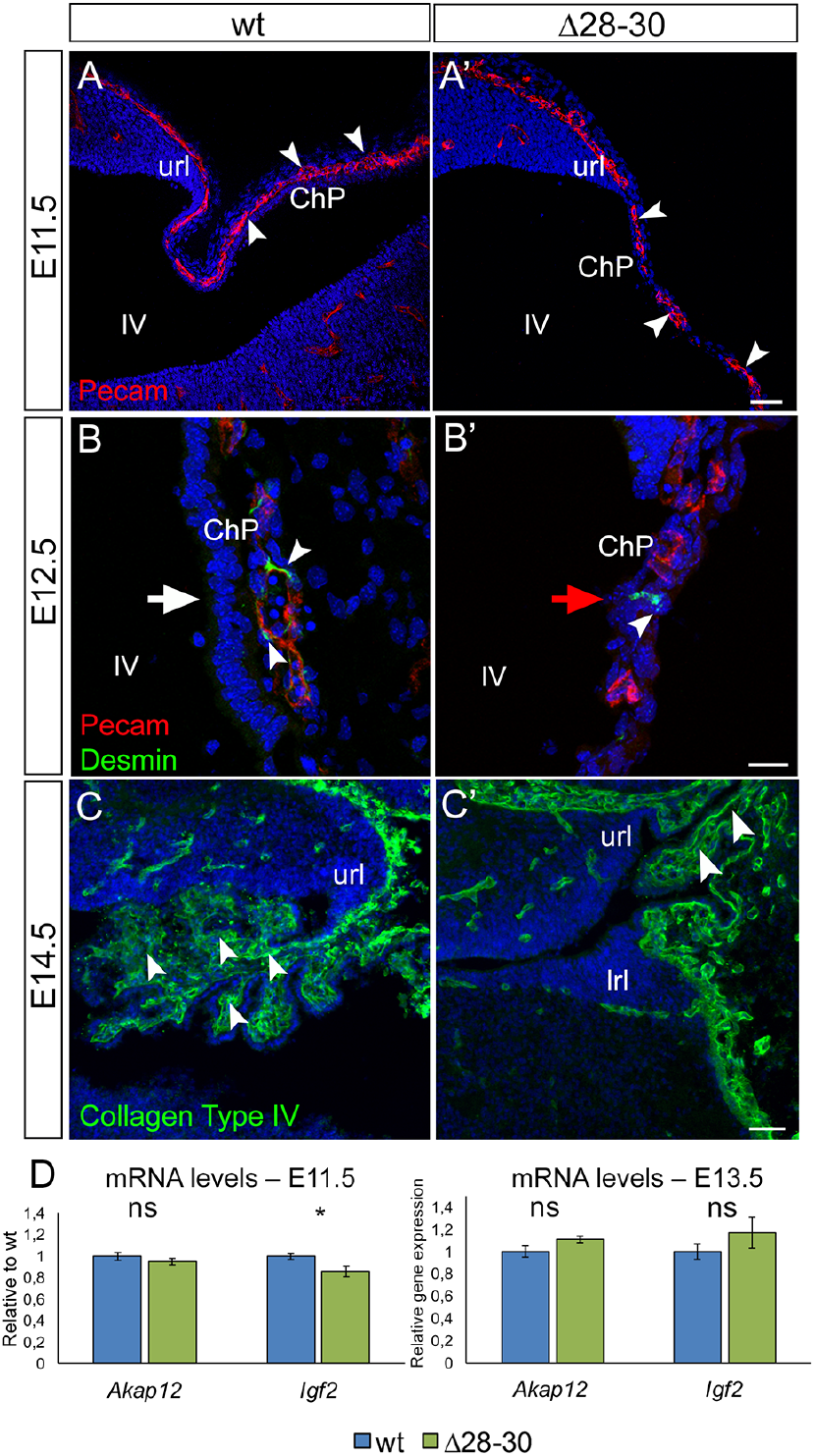
Vascular and pericyte markers are expressed in the mutant hChP. A-C’: parasagittal cerebellar sections harvested at different developmental stages and stained for different markers of the hChP stroma, vascular tree, and basement membrane. A,A’: Pecam-positive capillaries are present in the mutant hChP (white arrowheads in A and A’) at E11.5, albeit reduced in size. B,B’: E12.5 Desmin-positive pericytes are detected in mutant embryos. Note stratified epithelial nuclei in B (white arrow) and lack thereof in B’ (red arrowhead). C,C’: Collagen type IV is present in the basal membrane of the mutant hChP at E14.5. D: RT-qPCR reveal slight downregulation or unchanged expression of *Akap12* and *Igf2* in the mutant hindbrain at E11.5 (* p < 0.05,) and E13.5. ns = not significantly different. N=3/genotype; results are plotted as mean ± s.e.m is plotted; Welch’s unequal variances t-test. url: upper rhombic lip; lrl: lower rhombic lip; ChP: choroid plexus; IV: 4th ventricle. Size bar: 50μm in A,A’,C,C’; 20μm in B,B’.

### Changes in microvilli and tight junctions

As mentioned, ChP epithelial cells exhibit microvilli and cilia on their apical plasma membrane. Microvilli expand the luminal surface of the epithelial cell to increase the capacity of the ChP to secrete CSF. Ezrin is expressed in the ChP (Gimeno et al., 2004) and concentrates in the cytoskeleton of apical microvilli (Viswanatha et al., 2014). An ezrin antibody was used to decorate hChP microvilli on E12.5 parasagittal sections. Whereas the apical margin is strongly positive in the wt hChP epithelium (FIG. 6A, white arrowheads), ezrin signal is weaker, patchy and poorly polarized in the mutant epithelium, and it is also present in the basal domain of some hChP cells (Fig. 6B’, red arrowheads), pointing to a possible defect in cell polarity (see supplemental movies 1 and 2). Conversely, in the developing tChP (Suppl. Fig. 2A,B’), ezrin is expressed at high levels and apically located in both wt and mutant epithelia (white arrowheads). These results further confirm that the observed cytoarchitectural changes are highly specific to the hChP.

**Figure 6.**
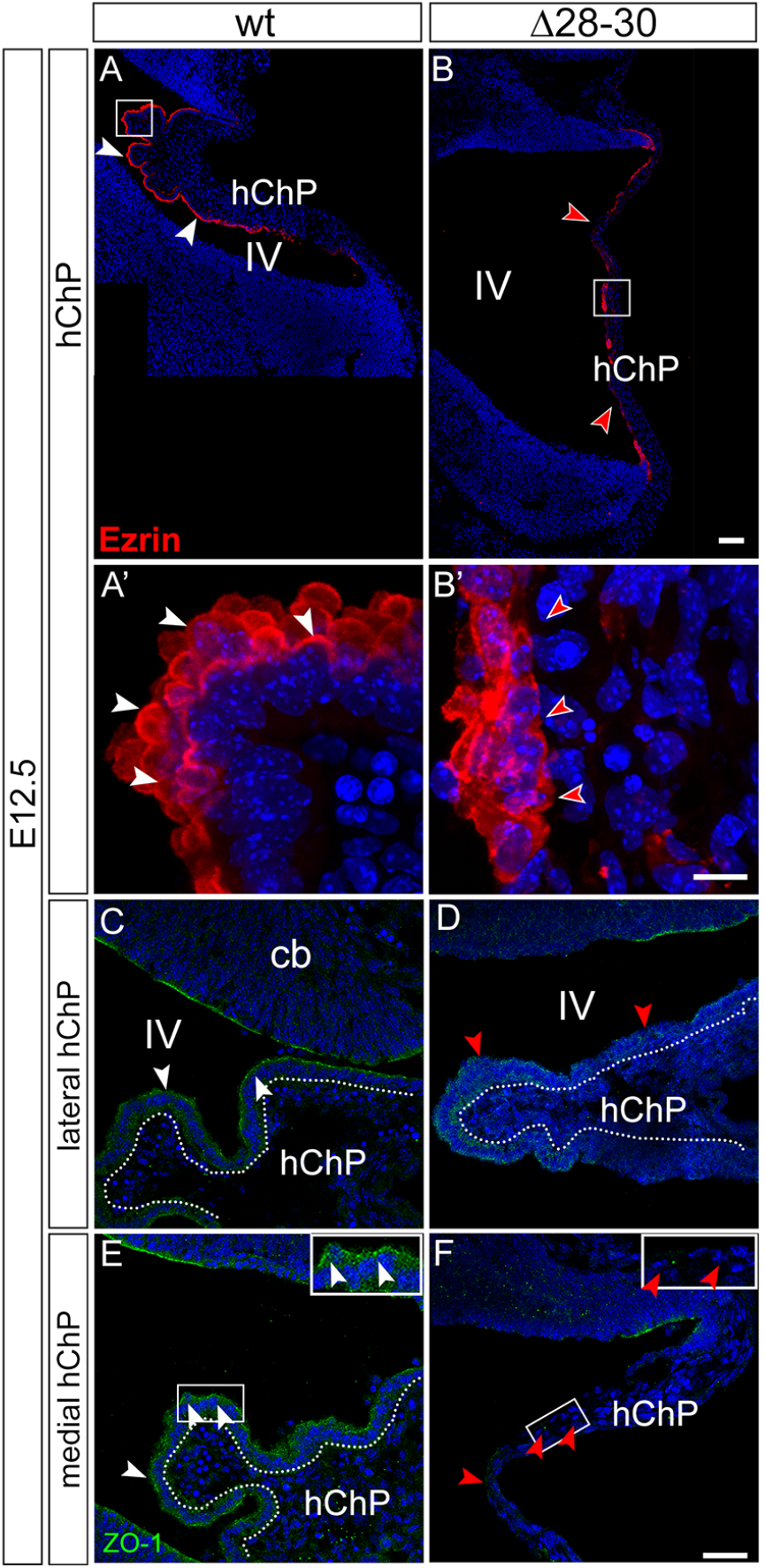
Expression of epithelial markers is downregulated in the hindbrain choroid plexus of *Zfp423* mutants. A-B’: parasagittal cerebellar sections at E12.5 stained for the apical microvilli marker Ezrin. The apical margin is strongly positive in the wt hChP epithelium (A,A’, white arrowheads); however, Ezrin signal is weaker, patchy and poorly polarized in the mutant epithelium (B,B’, red arrowheads), extending to the basal domain of some hChP cells (B’, red arrowheads). C,F: the signal of the tight junction marker ZO-1 is weaker and discontinuous in mutant lateral sections (red arrowheads in D) compared to wt, while it is virtually absent from the mutant hChP epithelium in medial sections (red arrowheads in F). Dotted lines in C,D,E highlight the columnar hChP epitelium, not visible in F. cb: cerebellum; hChP: hindbrain choroid plexus; IV: fourth ventricle. Size bar: 100μm in A, B; 10μm in A’, B’; 40μm in C-F.

Moreover, we performed an immunofluorescence with a zonula occludens 1 (ZO-1)- specific antibody to visualize the tight junctions that interconnect ChP epithelial cells at their apical pole. This marker is downregulated in medial and lateral sections of the Zfp423 mutant hChP at E12.5 (Fig. 6 D,F red arrowheads) compared to the wt (Fig. 6 C,E white arrowheads), suggesting that the apico-basal segregation of membrane proteins, normally maintained by tight junctions, may be compromised, possibly leading to a leakage of the blood-CSF barrier.

### Impaired generation of multiciliated cells in the mutant hChP

Specialized ependymal cells of the wt hChP are oligo-ciliated, exhibiting apical tufts of motile cilia (9 + 2 microtubules) ranging in number from 4 to 20/cell. These cilia express Arl13b in the axoneme while γ-tubulin labels the basal body (reviewed in Beisson and Wright, 2003; e.g. Casoni et al., 2017). Immunostaining with Arl13b and γTubulin Abs was used to characterize ciliogenesis in parasagittal hChP sections. At E11.5, while the wt hChP contains some multiciliated epithelial cells, and basal body duplication is evident (Fig. 7A,A’, white arrowheads), only monociliated cells are visible in the mutant hChP (Fig. 7B,B’, red arrowheads). Both lateral and medial parasagittal sections of wt E12.5 embryos display a columnar hChP epithelium characterized by numerous tufts of Arl13b+ cilia with multiple γTubulin+ basal bodies (Fig. 7C,E, insets in C’,E’, white arrowheads). Conversely, in mutant littermates the lateral sections display a smaller, cuboidal and disorganized hChP epithelium with poorly aligned γTubulin dots and absent, or very rare, Arl13b+ axonemes (Fig. 7D, inset in D’ red arrowheads). More strikingly, medial sections contain a monostratified and squamous hChP epithelium, and only sparse monociliated cells are present (Fig. 7F, inset in F’, red arrowheads). However, the analysis of the tChP in the lateral ventricles revealed that wt and mutant cells alike display a columnar and multiciliated epithelium positive for Arl13b+ and γTubulin (data not shown), again suggesting that *Zfp423* is only required for the specification of the hChP ciliated epithelium and that some alternative regulatory molecules may be in place in the embryonic lateral ventricles.

**Figure 7.**
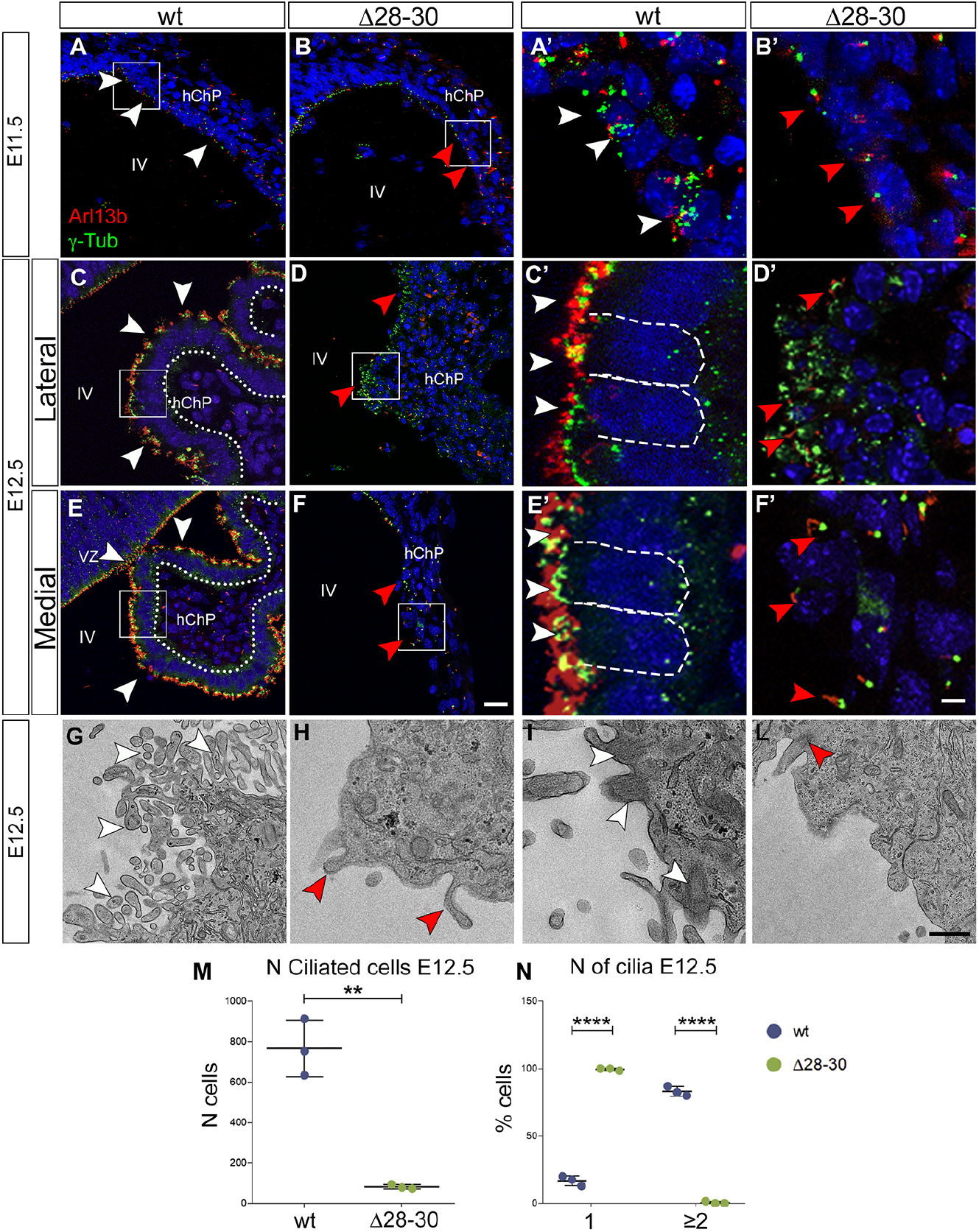
The Zfp423 mutant ChP fails to develop a multiciliated epithelium. A-F’: parasagittal cerebellar sections of the hChP were obtained at different developmental stages and stained for Arl13b and γTubulin, to visualize cilia and basal bodies, respectively. At E.11.5, wt hChP epithelial cells display numerous γTubulin-positive basal bodies and several Arl13b-positive cilia (A,A’ white arrowheads); conversely, the mutant hChP contains mono-ciliated cells (B,B’ red arrowheads). At this stage, both mutant and wt hChP display a cuboidal epithelium (white arrows). At E12.5, the wt epithelium is multiciliated, with several γTubulin-positive and Arl13b-positive dots on the apical domain of the epithelium, that are columnar in the wt (C,C’,E,E’, white arrowheads, dashed lines), both in lateral and in medial sections. Instead, the mutant hChP epithelium is disorganized, consisting of poorly aligned cuboidal cells with several γTubulin-positive dots but few and sparse Arl13b-positive cilia (D,D’, white arrowheads). Medial sections (F,F’) contain a monostratified and squamous hChP epithelium, and only sparse monociliated cells (red arrowheads). G-L: transmission electron microscopy images of the ChP epithelial cells at E12.5; wt epithelial cells display several microvilli on their apical surface (G, white arrowheads) and several cilia (I, white arrowheads); however, the mutant epithelium displays sparse microvilli (H, red arrowheads) and a single cilium, when present (L, red arrowhead). M. Graphs show the number of ciliated cells in wt and in mutant sections at E12.5. N=3/genotype. A two-tailed unpaired t test was performed to analyze differences between samples (**, p < 0.005). N. Graphs show the percentage of cells with 1 cilium or ≥2 cilia. N=3/genotype. Two-way ANOVA with Bonferroni post-hoc analysis (****, p < 0.0001). Results are plotted as mean ± SD. hChP: hindbrain choroid plexus, IV: 4th ventricle. Size bar: 20μm in A-F; 5μm in A’-F’; 1μm in G-L.

To achieve a better resolution, we analyzed the ultrastructure of the E12.5 hChP epithelium by transmission electron microscopy. Our results reveal that while wt epithelial cells contain numerous microvilli (Fig. 7G, white arrowheads) and are multiciliated (Fig. 7I, white arrowheads), mutant cells project few if any microvilli (Fig. 7H, red arrowheads) and, occasionally, a single cilium (Fig. 7L, red arrowheads).

At E12.5 the mutant hChP epithelium shows a sharp and significant decrease in the number of ciliated cells (Fig. 7M, N=3/genotype, ** p < 0.005). By counting cells with 1 cilium or with ≥ 2 cilia, we showed that the mutant hChP displays close to 100% monociliated cells and hardly any oligociliated cells (Fig. N, n=3/genotype, **** p < 0.00001).

Next, we analyzed the hChP at late gestation (E18.5) and postnatal stages (P18) by immunostaining frontal sections with Arl13b and γTubulin Abs. At E18.5, wt embryos exhibit an overwhelming majority of multiciliated cuboidal cells in the hChP (Supp Fig 3A,A’ white arrowheads), although sparse monociliated cells are also detected in the lateral segment of the hChP (Supp Fig 3A,A’ white arrow). However, mutant embryos only present a minute lateral segment of the hChP, close to the Luschka foramina (Supp Fig 3B,B’), with cuboidal monociliated cells (Supp Fig 3B,B’ white arrows) and some oligociliated cells (Supp Fig 3B,B’ white arrowheads). In this lateral territory, the mutant epithelium has fewer ciliated cells (Suppl. Fig. 3I, N=3/genotype, **, p < 0.005), and 20% less oligociliated cells than the wt (Suppl. Fig. 3L, n=3/genotype, **, p < 0.005).

The central segment of the hChP, normally observed in the 4th ventricle of wt embryos (Suppl. Fig. 3C,C’), was totally absent in the mutant (Suppl. Fig. 3D,D’). The same defects were observed at P18 (Suppl. Fig. 3E-H’).

### Zfp423 regulates genes critically involved in hindbrain patterning and multiciliogenesis

The molecular mechanisms involved in the development of the ChP epithelium have been partially characterized (Liddelow, 2015; Lun et al., 2015b). The hChP derives from the hindbrain roof plate. By wholemount in situ hybridization, we examined the expression of *Gdf7,* a highly specific RP marker (Lee et al., 1998), in E10.5 embryos (Fig. 8A). While *Gdf7* is expressed in the mutant RP, and its overall levels are unchanged, the signal is discontinuous flanking the fourth ventricle (red arrowheads in A’). By RNA sequencing (RNA-seq), we conducted a genome-wide transcriptome analysis (see materials and methods) on a 700 μm fragment of the E9.5 hindbrain spanning rhombomeres 1-6 harvested from *Zfp423* deletion mutants (Δ28-30, Casoni et al., 2017) and their wildtype littermate (results shown in supplemental Fig. 4). The results of our analysis show changes in the expression of regulatory genes critically involved in the patterning of the dorsal hindbrain and RP. In particular, our results show a sharp decrease in the levels of *Wnt1,* a strong upregulation of *Wnt3* and *Wnt8a,* whose role in this process has not been characterized to date, and a marked upregulation of the extracellular BMP signaling antagonist *Grem1,* accompanied by downregulation of the intracellular BMP target *Msx2.* These results were confirmed by RT-qPCR at E9.5 and E10.5 (Fig. 8B,C). Other differentially expressed genes identified in our transcriptome analysis are key players in multiciliogenesis: *Gmnc* (a.k.a. *GemC1*) is the master regulator of the transition from monociliated neurogenic stem cells to multiciliated, mitotically quiescent ependymal cells (Zhou et al., 2015). A dramatic decrease of *Gmnc* expression was scored in the mutant hindbrain. This result was validated by RT-qPCR (Fig. 8B,C): *Gmnc* is >95% reduced in E9.5 mutants (****, p < 0.00001; Fig. 8B) and 50% reduced at E11.5 (**, p < 0.005; Fig. 8C) and E13.5 (**, p < 0.005; data not shown). The results of in situ hybridization conducted at E10.5 showed that whereas the wt hChP epithelium is strongly positive for *Gmnc* in both lateral and medial sections (Fig. 8 D,E white arrowheads), in mutants *Gmnc* signal is weaker and patchy in the lateral hChP epithelium, and virtually absent in medial sections (Fig. 8 D’,E’ red arrowheads). *Gmnc*functions upstream of *Mcidas* (a.k.a. *Linkeas* or *multicilin*) (Spassky and Meunier, 2017; Zhou et al., 2015). RT-qPCR conducted in the E11.5 hindbrain (Fig. 8C) showed that *Mcidas* is significantly downregulated in the mutant (**, p < 0.005). Likewise, *Foxj1,* a marker of multiciliated epithelia, encoding a regulator of multiciliated gene transcription and of basal body docking to the apical cytoskeleton (Gomperts et al., 2004), is also significantly downregulated at E11.5 (Fig. 8C, p < 0.001). Taken together, these results point to an overall deregulation of multiciliated cell fate determination.

**Figure 8.**
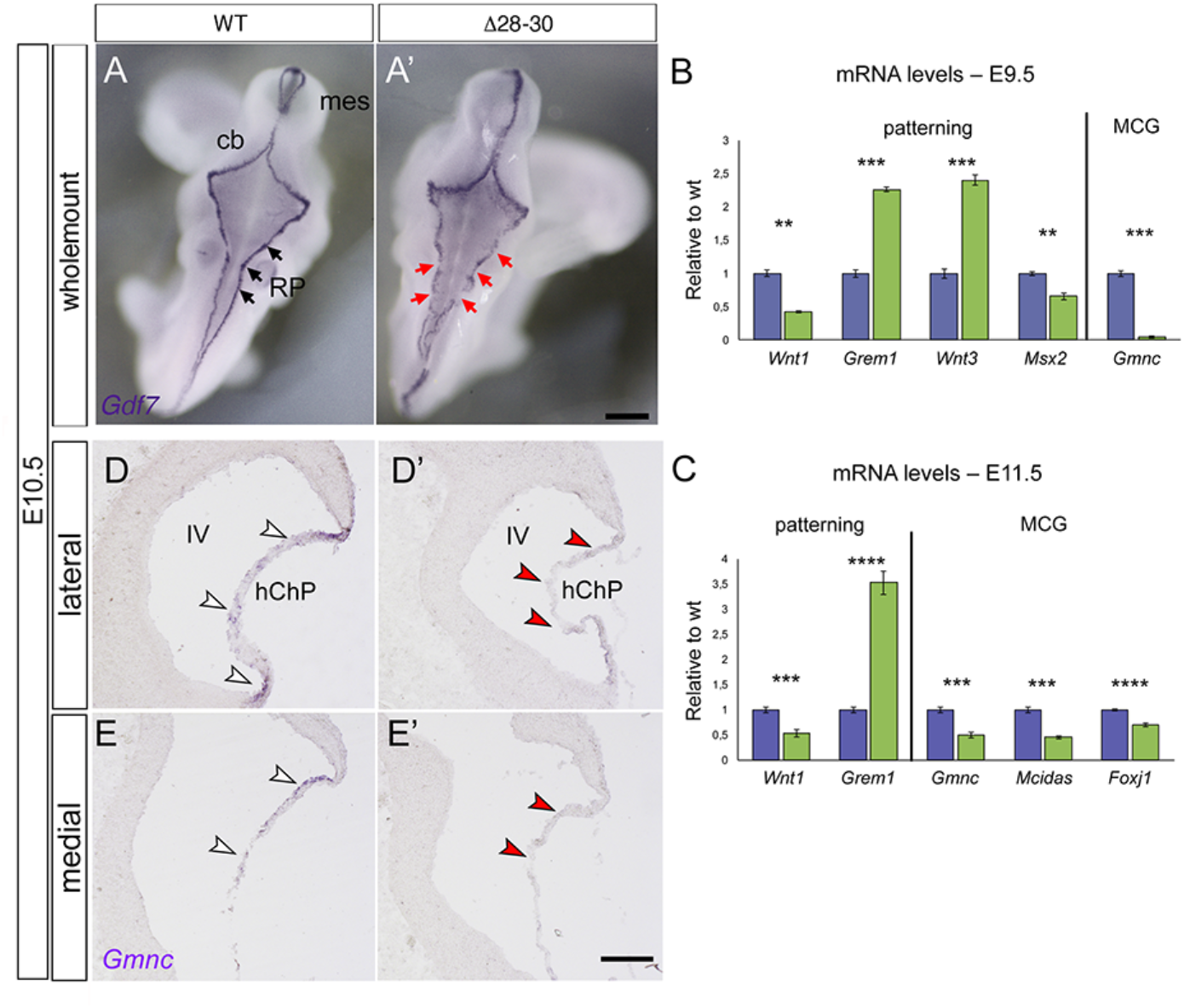
The mutant hChP shows changes in markers of roof plate patterning and ependymal cell specification. A: Dorsal view of wholemount *in situ* hybridization reveals a conserved but discontinuous *Gdf7* expression pattern in the mutant (red arrows). B: RT-qPCR quantitation of roof plate patterning markers at E9.5 reveals deregulated expression of four early patterning molecules involved in Wnt and BMP signaling and a profound downregulation of the multiciliogenesis master gene *Gmnc.* C: deregulation of *Wnt1* and *Grem1* persists at E11.5, together with downregulation of three key regulators of multiciliogenesis (see text). D-E’: *in situ* hybridization of hChP E10.5 parasagittal sections shows discontinuous expression of *Gmnc* in lateral sections of the mutant hChP compared to the wt (D,D’), and complete lack of the transcript in medial *Zfp423*-mutant sections (E,E’). MCG: multiciliogenesis markers. cb: cerebellum, mes: mesencephalon, hChP: hindbrain choroid plexus, IV: 4th ventricle. All values are expressed mean ± s.e.m. (C, n= 3/genotype; p < 0.001; Welch’s t-test), at E11.5 (D, n= 3/genotype; p < 0.001; Welch’s t-test). Size bar: 500μm in A,A’; 100μm in D-E’.

## DISCUSSION

In the present paper, we examine the effects of *Zfp423* mutation on hChP development. We find that the hChP is completely absent in homozygous mutants, except for a minute symmetrical vestige of this structure restricted to the lateralmost region of the fourth ventricle, right next to the foramina of Luschka. During early embryonic development, the mutant hChP is misfolded, contains very few recognizable ependymal cells and is functionally compromised, failing to express the thyroid hormone and retinoic acid transporter transthyretin, and the water transporter aquaporin 1. Moreover, the mutant hChP loses the expression of two established markers of the embryonic hChP that are essential for its development: *Lmx1a* and *Otx2* (Johansson et al., 2013; Millonig et al., 2000).

Fate-mapping studies have established that the hChP emerges from two postmitotic morphogenetic fields, medial and lateral, both double-positive for *Wnt1* and *Gdf7* (Hunter and Dymecki, 2007). Along the antero-posterior axis, the hChP is thought to develop mainly from a transition zone adjacent to the LRL, expanding caudally from a rostral bud of newly formed ependymal cells. The caudal part of the hChP derives from rhombomeres 2–8, whereas the more rostral one derives from rhombomere 1 (Hunter and Dymecki, 2007). However, the early expression domains of some key genes in the patterning of this territory, including *Wnt1, Gdf7, Lmx1a* and *Otx2,* span the entire longitudinal expanse of the 4th ventricle. Similarly, the expression domains of genes involved in the differentiation of multiciliated ependymal cells (e.g. *Gmnc* and *Foxj1)* also extend throughout the length of the 4th ventricle. Likewise, ZFP423 protein expression spans URL and LRL starting at E9.5.

### Zfp423 controls the earliest stages of hChP development

The results of studies conducted on murine and avian models clearly indicate that ChP patterning and cell fate specification starts early in development. In the mouse brain, hChP fate is determined starting between E8.5 and E9.5 (Thomas and Dziadek, 1993), 2–3 days before the hChP primordium becomes anatomically recognizable. Likewise, chicken and quail grafting studies have shown that ChP fate is determined up to 3 days before its anatomical appearance (Wilting and Christ, 1989).

*Zfp423* expression initiates in the hindbrain at the open neural plate stage (E7.5) and transcript levels increase between E8.5 and E9.5, with the transcript spanning the upper and lower rhombic lip, and the roof plate (our unpublished results). Because of its precocious expression in the prospective hindbrain, starting at the open neurula stage, ZFP423 controls very early and relatively unexplored stages of hChP development. In the mouse, those stages precede by at least two days the beginning of SHH-dependent progenitor proliferation that controls the expansion of the hChP from its earliest primordium (described in Huang et al., 2009b).

### Zfp423 is required for hChP ependyma development

Choroid plexuses develop thanks to the interaction between the neuroepithelium, which gives rise to the multiciliated ependyma, and mesodermic derivatives, including the prospective meninges, pericytes and fenestrated blood vessels.

Sophisticated experiments conducted in chick embryos (Broom et al., 2012) have led to the conclusion that the ectoderm-derived roof plate boundary organizer, located caudal to the rhombic lip and strongly positive for *Zfp423,* signals to the roof plate itself to specify the expression of early choroid plexus markers. Later, SHH-expressing hChP ependyma plays an instructive role on both adjacent epithelial progenitors and underlying vascular outgrowth (Huang et al., 2009b; Nielsen and Dymecki, 2010).

Because the ZFP423 protein is strongly expressed in both epithelial and mesenchymal constituents of the developing hChP, we analyzed the impact of *Zfp423* mutation on several cell types populating the hChP. ZFP423 decorates the prospective leptomeningeal mesenchyme at very early stages of RP development (E11.5), prior to the appearance of a recognizable ChP. However, in the mutant, early leptomeningeal markers are conserved or only partially downregulated. Likewise, the expression of endothelium and pericyte markers is maintained, albeit slightly decreased, and *Zfp423* mutation strongly affects the epithelial component, which fails to express the defining features of the ChP ependyma. In fact, the mutant epithelium evolves into a monolayer of squamous cells, rather than giving rise to a columnar cell layer first and a ciliated cuboidal monolayer later.

### Transcriptome profiling points to roles for Zfp423 in hindbrain patterning

Several factors whose expression is significantly decreased in the mutant are critically involved in hindbrain patterning, and include Wnt and BMP signaling molecules. Wnt signaling is a key factor in ChP development, both in the telencephalon and rhombencephalon (inter alia Awatramani et al., 2003; Grove et al., 1998). In the *Zfp423-mutant* hChP primordium, *Wnt1* is sharply downregulated starting at E9.5, while other canonical Wnt ligand genes are significantly upregulated, as in the case of *Wnt3* (Fig. 8) and *Wnt8a* (not shown), whose roles in Chp development have not been addressed. No significant changes were observed at those early stages in the levels of *Wnt5a,* encoding a Wnt planar polarity molecule that, slightly later in development, is critically involved in hindbrain morphogenesis (Kaiser et al., 2019). In the hChP epithelium, *Wnt1* is co-expressed with *Gdf7,* encoding a RP-specific member of the BMP protein family. Conditional disruption of the *BMP receptor 1* gene in the telencephalon severely impairs tChP development (Hebert et al., 2002). Moreover, in vitro studies have revealed that BMP signaling is sufficient to convert ESC-derived neuroepithelium into choroid plexus epithelial cells (Watanabe et al., 2012). While the *Gdf7* transcript is not reduced quantitatively in the mutant hChP, its expression domain is discontinuous and reveals alterations in RP morphogenesis. Likewise, BMP signaling is perturbed in the mutant, with a sharp upregulation of *Grem1,* an extracellular BMP antagonist belonging to the gene family that comprises the head-inducing factor Cerberus and the tumor suppressor DAN (Hsu et al., 1998). Accordingly, the mutant exhibits a significant downregulation of the SMAD1-SMAD4 transcriptional target *Msx2.* These results suggest a hierarchically elevated role for *Zfp423* within the regulatory cascades that govern dorsal hindbrain patterning.

### Zfp423 is required to activate multiciliogenesis in the hChP

In addition to the deregulation of signaling pathways controlling hindbrain patterning, *Zfp423* mutants also feature a highly significant downregulation of genes promoting multiciliogenesis. Our RNA-seq analysis has revealed that, in the mutant, *Gmnc* expression is totally absent at E9.5, corresponding to the onset of hChP morphogenesis. *Gmnc* mediates the conversion of proliferating, monociliated neuroepithelium into postmitotic, multiciliated ChP ependymal cells (Arbi et al., 2016; Lalioti et al., 2019), uncoupling basal body proliferation from cell division. *Gmnc* controls the expression of other factors involved in the same process, namely *Mcidas* (a.k.a *Linkeas)* and *Foxj1* (Lalioti et al., 2019), which are also downregulated. While most differentially expressed genes are upregulated in the mutant (and this includes *Zfp423* itself), confirming the notion that ZFP423 acts as a transcriptional repressor (Cho et al., 2013), *Gmnc* transcription is drastically repressed in the mutant hindbrain. This finding suggests that *Gmnc* may be an indirect target of ZFP423.

### Zfp423 is not required for tChP development

In *Zfp423* mutants, the telencephalic ChP, located in the prospective lateral ventricles, develops rather normally in terms of gross appearance and cytoarchitecture, consistent with the finding that rhombencephalic and telencephalic choroid plexuses differ greatly in their positional identities and secretomes (Lun et al., 2015a). *Zfp423,* which affects telencephalic midline induction (Cheng et al., 2007) and neocortical neurogenesis (Massimino et al., 2018), may not be absolutely required for patterning or multiciliated cell specification in the telencephalic midline, suggesting that an unidentified functional homolog operates at that site.

### Conclusions

In summary, *Zfp423* is a master gene and one of the earliest known determinants of hChP development. Its mutation severely impairs hChP morphogenesis and the expression of key regulatory signals required for its development and function. The defective production and/or secretion of key molecules by the hChP may be one causal factor of the overall phenotype observed in *Zfp423* mutant mice and possibly contribute to the pathogenesis of cerebellar vermis hypoplasia and Joubert-like cerebellar malformations in patients carrying *ZNF423* mutations (Chaki et al., 2012).

## Acknowledgements

Image analysis was carried out at ALEMBIC, an advanced microscopy laboratory established by the San Raffaele Scientific Institute and Vita-Salute San Raffaele University. Library preparation and RNA sequencing was conducted at the CTGB (Center for Translational Genomics and Bioinformatics, San Raffaele Scientific Institute). We thank F. Marroni and A. Pisciottani for critical discussion of the results and manuscript, and Angelo Quattrini for help with transmission electron microscopy. Special thanks to Søren Warming for making the Δ28-30 line available to us.

## Author contributions

Conceptualization: F.C., L.C., O.C., G.G.C.; methodology: F.C., L.C., F.V., P.P., G.G.C.; formal analysis: F.C., L.C., G.G.C.; investigation: F.C., L.C., F.V., P.P.; writing - original draft: F.C., L.C., G.G.C.; writing - review & editing: F.C., L.C., G.G.C.; Bioinformatic analysis: L.M.; Supervision: F.C. and G.G.C.; Project administration: F.C.; Funding acquisition: G.G.C.

## SUPPLEMENTAL MATERIALS AND METHODS

### Primer sequences

Genotyping primers:

*Δ28-30*-wtF: 5’-TGAGAGAGGACACCTACTCT-3’;

*Δ28-30*-wtR: 5’-GCAGGGAGCAAACTGTCTCTT-3’;

*Δ28-30*-mutR: 5’-GGTGTGACCTTTGTGCGAGA-3’.

RT-qPCR primers:

*Ttr* F: 5’-AAAGTCCTGGATGCTGTCCG-3’

*Ttr* R: 5’-TTCTCATCTGTGGTGAGCCC-3’

*Aqp1* F: 5’-CAGTACCAGCTGCAGAGTGC-3’

*Aqp1* R: 5’-CATCACCTCCTCCCTAGTCG-3’

*Lmx1a* F: 5’-TGAGTGTCCGTGTGGTTCAG-3’

*Lmx1a* R: 5’-CCCGCATTCCCACTACCATT-3’

*Otx2* F: 5’-CTGACCTCCATTCTGCTGCT-3’

*Otx2* R: 5’-GGAAGAGGTGGCACTGAAAA-3’

*Gmnc* F: 5’-GAGGCTCAGCTCTCATCTCA-3’

*Gmnc* R: 5’-ATGACAGCAACTTCTTGGCC-3’

*Mcidas* F: 5’-CCAGCTCTCACAACCATAGAC-3’

*Mcidas* R: 5’-GCATCTCTGAAATTCTGCAGG-3’

*Msx2* F: 5’-GGAAAATTCCGAAGACGGAG-3’

*Msx2* R: 5’-CTTCCGGTTGGTCTTGTGTT-3’

*Grem1* F: 5’-CCTTTCTTTTTCCCCTCAGC-3’

*Grem1* R: 5’-ACAGCGAAGAACCTGAGGAC-3’

*Wnt1* F: 5’-AAATGGCAATTCCGAAACC-3’

*Wnt1* R: 5’-GAGGTGATTGCGAAGATGAA-3’

*Wnt3* F: 5’-CTTCTAATGGAGCCCCACCT-3’

*Wnt3* R: 5’-GAGGCCAGAGATGTGTACTGC-3’

*Gdf7* F: 5’-GGCTTCACAGACCAAGCAAC-3’

*Gdf7* R: 5’-GCACTGTCCCTGTCTGGTTC-3’

I*gf2* F: 5’-GGTTTGCATACCCGCAGCA-3’

I*gf2* R: 5’-CACAAGGCGAAGGCCAAAGA-3’

*Akap12* F: 5’-CCTGACAGAATCCTAAGACGTG-3’

*Akap12* R: 5’-GGTTGAAATCATTGGACGGC

*Foxj1* F: 5’-ACGGACAACTTCTGCTACTTC-3’

*Foxj1* R: 5’-CTCCCGAGGCACTTTGATG-3’

*β-actin* F: 5’-CTGTCGAGTCGCGTCCACC-3’

*β-actin* R: 5’-TCGTCATCCATGGCGAACTG-3’

