## Supplemental figures for "ZFP423 regulates early patterning and multiciliogenesis in the hindbrain choroid plexus"

### SUPPLEMENTAL FIGURE LEGENDS

#### Supplemental Figure 1

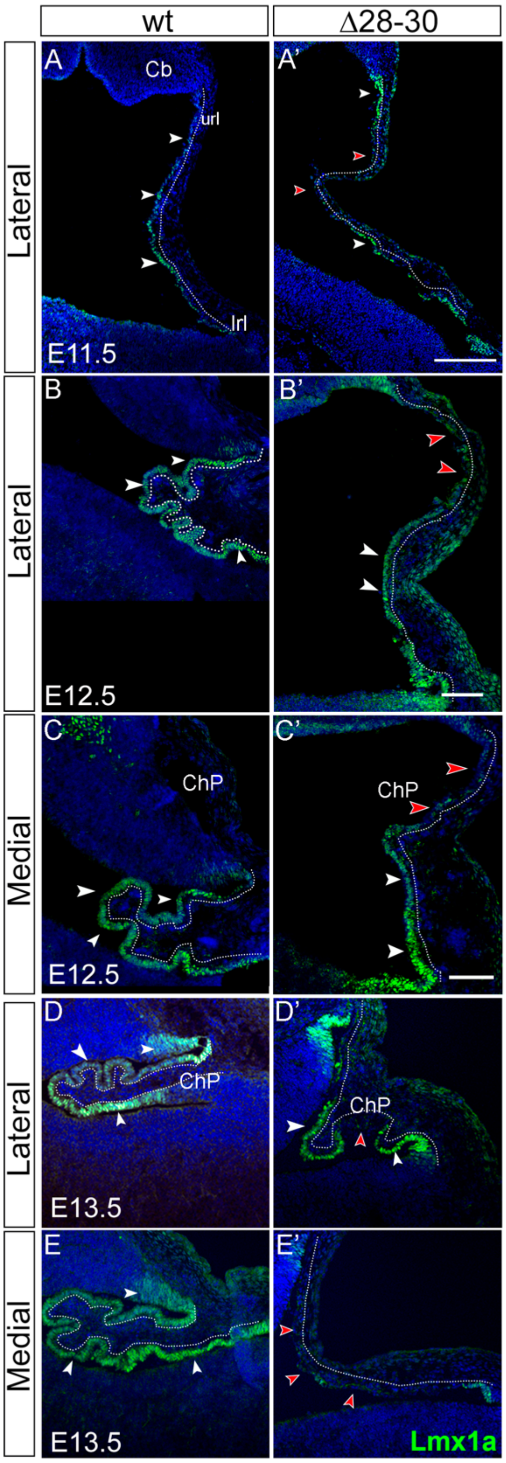


**Loss of Lmx1a protein expression in the *Zfp423* mutant epithelium.** A-E': parasagittal cerebellar sections at different stages of development, stained for Lmx1a to visualize its distribution in the developing hChP. A,A' Lmx1a immunostaining of E11.5 sagittal sections show a discontius decoration of this protein on the mutant epithelium. B-C', lateral and medial parasagittal cerebellar sections at E12.5 show an increase in the expression of the protein in the wt columnar epithelium (white arrowheads and dotted lines in C and D), while the expression in mutant epithelium is discontinuous (red arrowheads and dotted lines in C' and D'). E,E', medial parasagittal cerebellar sections at E13.5 show an increase in the expression of the protein in the wt columnar epithelium (white arrowheads and dotted lines in E), while the expression in mutant epithelium is absent (red arrowheads and dotted lines in E').

**Supplemental figure 2**

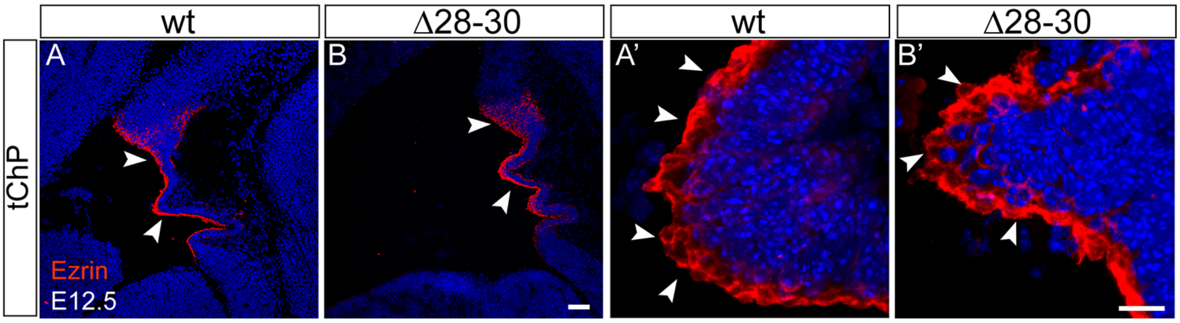


**The apical microvilli marker Ezrin is normally expressed in the mutant tChP.** A-B': Parasagittal sections of the lateral ventricles at E12.5, stained fort Ezrin. Ezrin is expressed at high levels and apically located in wt and mutant epithelia alike in the developing tChP (white arrowheads). Size bar: 100μm in A, B; 10μm in A', B'.

### Supplemental figure 3

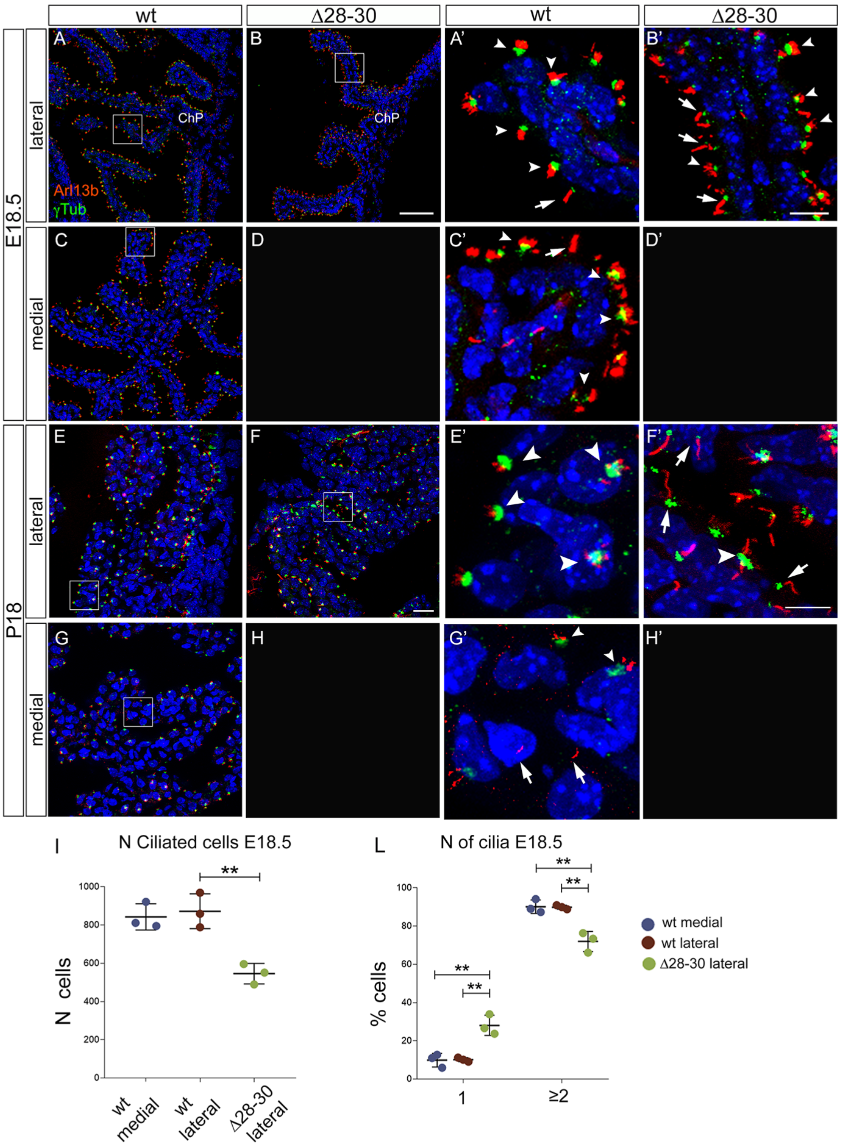


**Ciliary defects persist in the mutant hChP at birth and in adulthood.** A-H': parasagittal cerebellar sections at late embryonic (E18.5) and postnatal (P18) stages, stained for Arl13b and γTubulin, to visualize cilia and basal bodies, respectively. At E18.5, most wt hChP epithelial cells are multiciliated, with multiple γTubulin+ basal bodies and Arl13b-positive cilia (A,A' white arrowheads), while few monociliated cells (A,A' white arrow) can be observed in lateral parasagittal sections of the hChP. Conversely, in the mutant, the minute hChP segment confined to the lateral most aspect of the 4th ventricle, exhibits more monociliated (C', white arrows) and fewer multiciliated (C', white arrowheads) cells than the wt. At E18.5, the medial segment of the hChP is totally absent in mutant embryos (D). At P18, the wt hChP epithelial cells display mostly multiciliated cells with multiple γTubulin-positive basal bodies and Arl13b-positive cilia (E,E'' white arrowheads), in lateral sections of the hChP. Conversely the small lateral segment of the mutant hChP exhibits mostly monociliated (F,F', white arrows) and only a minority of multiciliated (F,F' white arrowheads) cells compared to the wt. Again, the bulk of the hChP is deleted in mutant embryos (H). I: the histogram shows the number of ciliated cells in the lateral and medial section of the wt hChP, and in the lateral section of the mutant hChP at E18.5. Data are plotted as mean ± SD. N=3/genotype. A two-tailed unpaired (Welch's) t-test was performed to determine statistically significant differences between lateral sections of the wt and mutant (**, P < 0.005). L: The histogram the percentage of cells with 1 cilium or ≥2 cilia for genotype. A 2way ANOVA was performed to determine statistically significant differences between samples. Data are plotted as mean ± SD is plotted. Bonferroni post-hoc analysis was applied. (****, P < 0.0001). Size bar: 20μm in A,B,C; 10μm in A',B',C'; 20μm in E,F,G; 10μm in E',F',G'.

**Supplemental figure 4**

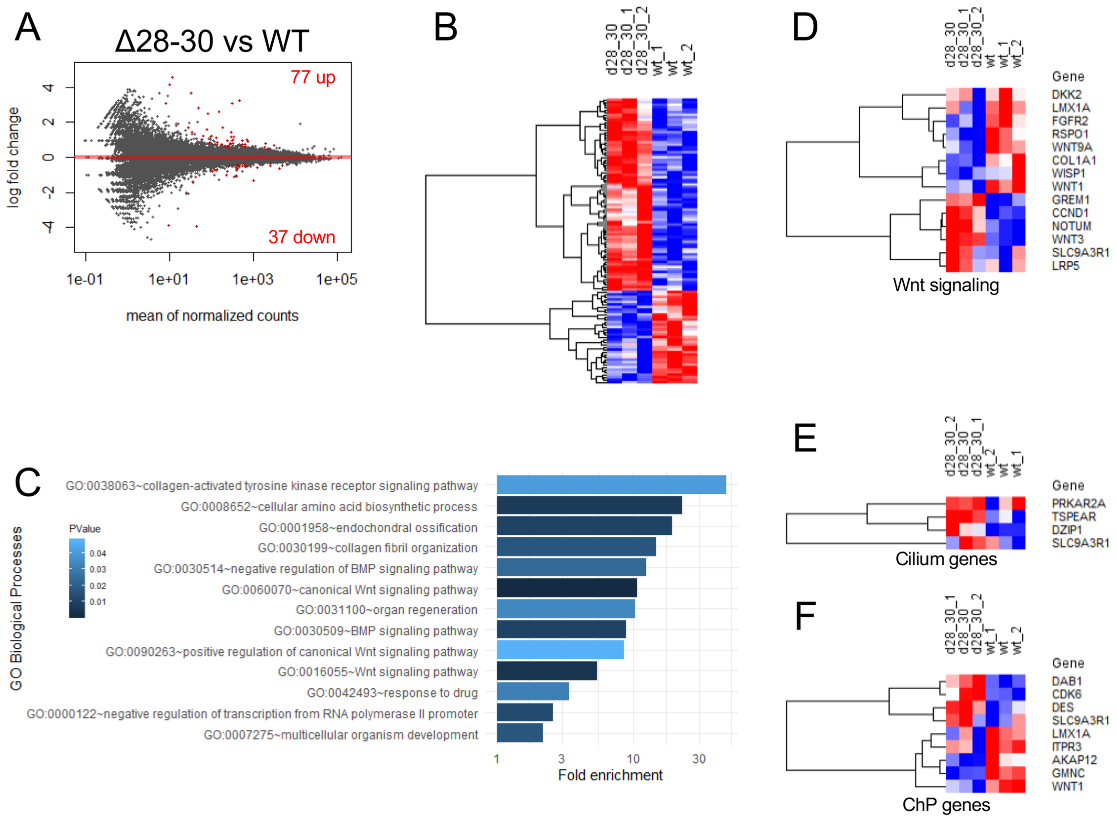


**Differential RNA sequencing analysis of wt vs Δ28-30 hindbrain at 9.5 days of embryonic development.** A. MA plot showing log2 fold change as a function of average gene expression. Differentially expressed genes (DEGs) with false discovery rate (FDR)<0.1 are highlighted in red. B. Heatmap of the DEGs with FDR<0.1. C. Functional enrichment analysis of gene ontology biological processes. D-F: Heatmaps showing DEGs related to Wnt signaling pathway (D), Cilium-associated (E), and Choroid plexus-associated (F) DEGs.
